# Biochemical analysis of the endoribonuclease activity of the human mitochondrial topoisomerase 1

**DOI:** 10.1101/2024.03.15.585299

**Authors:** Cyrielle P. J. Bader, Erika Kasho, Josefin M. E. Forslund, Malgorzata Wessels, Paulina H. Wanrooij

**Affiliations:** Department of Medical Biochemistry and Biophysics, Umeå University, 901 87 Umeå, Sweden

**Author notes:** These authors contributed equally (Cyrielle P. J. Bader, Erika Kasho).

## Abstract

The incorporation of ribonucleotides (rNMPs) into the nuclear genome leads to severe genomic instability, including strand breaks and short 2-5 bp deletions at repetitive sequences. Curiously, the detrimental effects of rNMPs are not observed for the human mitochondrial genome (mtDNA) that typically contains several rNMPs per molecule. Given that the nuclear genome instability phenotype is dependent on the activity of the nuclear topoisomerase 1 enzyme (hTop1), and mammalian mitochondria contain a distinct topoisomerase 1 paralog (hTop1mt), we hypothesized that the differential effects of rNMPs on the two genomes may reflect differing properties of the two cellular topoisomerase 1 enzymes. Here, we characterized the endoribonuclease activity of hTop1mt and found it to be less efficient than that of its nuclear counterpart, a finding that was partly explained by its substrate binding properties. While hTop1 and yeast Top1 showed higher affinity for an rNMP-containing substrate and were able to cleave at an rNMP located outside of the consensus cleavage site, hTop1mt showed no preference for rNMPs. As a consequence, hTop1mt was inefficient at producing the short rNMP-dependent deletions that are characteristic of Top1-driven genome instability. These findings help explain the tolerance of rNMPs in the mitochondrial genome.

## Introduction

The cellular concentrations of free ribonucleotides (rNTPs) clearly exceed those of deoxyribonucleotides (dNTPs), and single ribonucleotides are therefore occasionally inserted into the nascent DNA strand during DNA replication (Traut 1994; Kong et al. 2018; Nick McElhinny et al. 2010). Although the incorporated ribonucleotides (rNMPs) may have certain positive implications for genome stability such as facilitating some repair processes or relieving torsional stress, the net effects of rNMP incorporation into DNA are negative (Lujan et al. 2013; Cerritelli and Crouch 2016; Potenski and Klein 2014). Even single rNMPs jeopardize genome stability because their reactive 2’-hydroxyl group can induce single-stranded DNA breaks (Li and Breaker 1999). In addition, incorporated rNMPs alter the local structure and elasticity of the DNA and are thus expected to impact protein-DNA interactions (Jaishree et al. 1993; Chiu et al. 2014; DeRose et al. 2012).

To avoid these harmful consequences, rNMPs incorporated during nuclear DNA replication are efficiently removed by the ribonucleotide excision repair (RER) pathway that is initiated by RNase H2 (Reijns et al. 2012; Hiller et al. 2012; Rydberg and Game 2002; Sparks et al. 2012). The importance of rNMP removal is highlighted by the fact that mouse cells deficient in RNase H2 exhibit extensive genome instability in the form of increased DNA breaks, micronuclei and an activated DNA damage response (Reijns et al. 2012; Hiller et al. 2012). In the absence of RER, a lower level of rNMP removal can be mediated by the endoribonuclease activity of topoisomerase 1 (Top1) (Sekiguchi and Shuman 1997; Kim et al. 2011). However, Top1-mediated rNMP repair generates a nick flanked by a 2’,3’-cyclic phosphate that requires further processing prior to religation (Cho et al. 2013). At repetitive sequences where strand slippage can occur, Top1-mediated rNMP repair yields a distinct mutational signature consisting of 2-5 bp deletions (Kim et al. 2011; Sparks and Burgers 2015; Sekiguchi and Shuman 1997). Accordingly, the frequency of these 2-5 bp deletions as well as many of the other genome instability phenotypes of RNase H2-deficient cells are suppressed by the perturbation of *TOP1* in both yeast and human cells (Kim et al. 2011; Williams et al. 2013; Zimmermann et al. 2018).

RNase H2 and thus RER are absent from the mitochondrial compartment, whereby rNMPs incorporated during the replication of the mitochondrial genome (mtDNA) are not efficiently repaired but instead persist in this small multi-copy genome (Wanrooij et al. 2017; Wanrooij et al. 2020; Berglund et al. 2017; Moss et al. 2017; Grossman, Watson, and Vinograd 1973; Miyaki, Koide, and Ono 1973; Wong-Staal, Mendelsohn, and Goulian 1973). Based on our knowledge of rNMP biology in the nuclear genome, this absence of RER would be expected to predispose the mtDNA to rNMP-dependent instability (Wanrooij and Chabes 2019). Yet, there have to our knowledge been no reports of short rNMP-dependent deletions in mtDNA. Moreover, our previous work found no beneficial effects of decreasing the mtDNA rNMP load in mice *in vivo*, suggesting that the physiological level of rNMPs in mtDNA is well-tolerated (Wanrooij et al. 2020). Given that many of the negative effects of rNMPs on the nuclear genome are associated with their removal via the Top1-mediated repair pathway, we hypothesized that the tolerance of rNMPs in the mitochondrial genome may derive from different propensities of Top1-dependent ribonucleotide removal in the two cellular compartments.

In addition to the Top1 found in the nucleus, vertebrates contain a second Top1 paralog that is restricted to the mitochondrial compartment, Top1mt (Zhang et al. 2001). While Top1mt is not essential, *TOP1MT*^*-/-*^ mice exhibit an increase in negatively supercoiled mtDNA and Top1mt-deficient MEFs manifest with a defect in mitochondrial respiration (Zhang et al. 2014; Douarre et al. 2012). Human Top1mt (hTop1mt) and the human nuclear Top1 (hTop1) share 52% overall identity, with over 70% identity over the core and C-terminal domains that are the ones required for activity (Zhang et al. 2001; Stewart, Ireton, and Champoux 1997). Both hTop1mt and hTop1 belong to the Type 1B family of topoisomerases that can relax both negative and positive supercoils. Eukaryotic Top1s show strong preference for cleavage following the 3’ thymidine in the consensus sequence motif of 5’-(A/T) (G/C) (A/T) **T**^**↓**^-3’ (Champoux 2001). Their reaction mechanism involves the nicking of one strand in a double-stranded DNA substrate to form a covalent intermediate — termed the Top1-cleavage complex (Top1-cc) — between the active site tyrosine of the enzyme and the 3’ thymidine at the nick site (Champoux 2001). After rotating the substrate to relieve torsional strain, the enzymes catalyze re-ligation of the nick, consequently becoming released from the covalent complex (Champoux 2001). The reaction can be trapped in the Top1-cc stage using topoisomerase poisons like camptothecin (CPT) that inhibit the re-ligation step (Pommier et al. 2010).

When Top1 cleavage occurs at an embedded rNMP, the 2’-hydroxyl group of the rNMP can attack the Top1-cc phospho-tyrosyl bond to generate a 2’,3’-cyclic phosphate and liberate the enzyme prior to re-ligation (Sekiguchi and Shuman 1997) (Fig. 1a). This results in a nick flanked by a 2’,3’-cyclic phosphate and a 5’-hydroxyl group that cannot be ligated without further processing. Meticulous work on the *Saccharomyces cerevisiae* Top1 (scTop1) has revealed that the 2’,3’-cyclic phosphate is a preferred substrate of the enzyme and is rapidly turned over via one of two possible mechanisms: either scTop1 reverses the cyclization to re-generate the original rNMP-containing substrate, or it cleaves two nucleotides upstream of the nick to release a dinucleotide containing the 2’,3’-cyclic phosphate, generating a short gap (Sparks and Burgers 2015). Enzymatic removal of the Top1-cc from this second, upstream cleavage event and filling of the short gap results in an intact, rNMP-free DNA molecule (Fig. 1a, *green background*). Alternatively, at repetitive sequences that allow strand slippage to occur, the two DNA ends flanking the gap can be brought into close enough proximity for scTop1 to mediate their ligation (Fig. 1a, *red background*). Due to the strand re-alignment, this pathway leads to loss of one of the repeats in the repetitive sequence, explaining the characteristic signature of Top1-dependent 2-5 nt deletions at repetitive sequences (Sparks and Burgers 2015; Kim et al. 2011; Sekiguchi and Shuman 1997).

**Figure 1.**
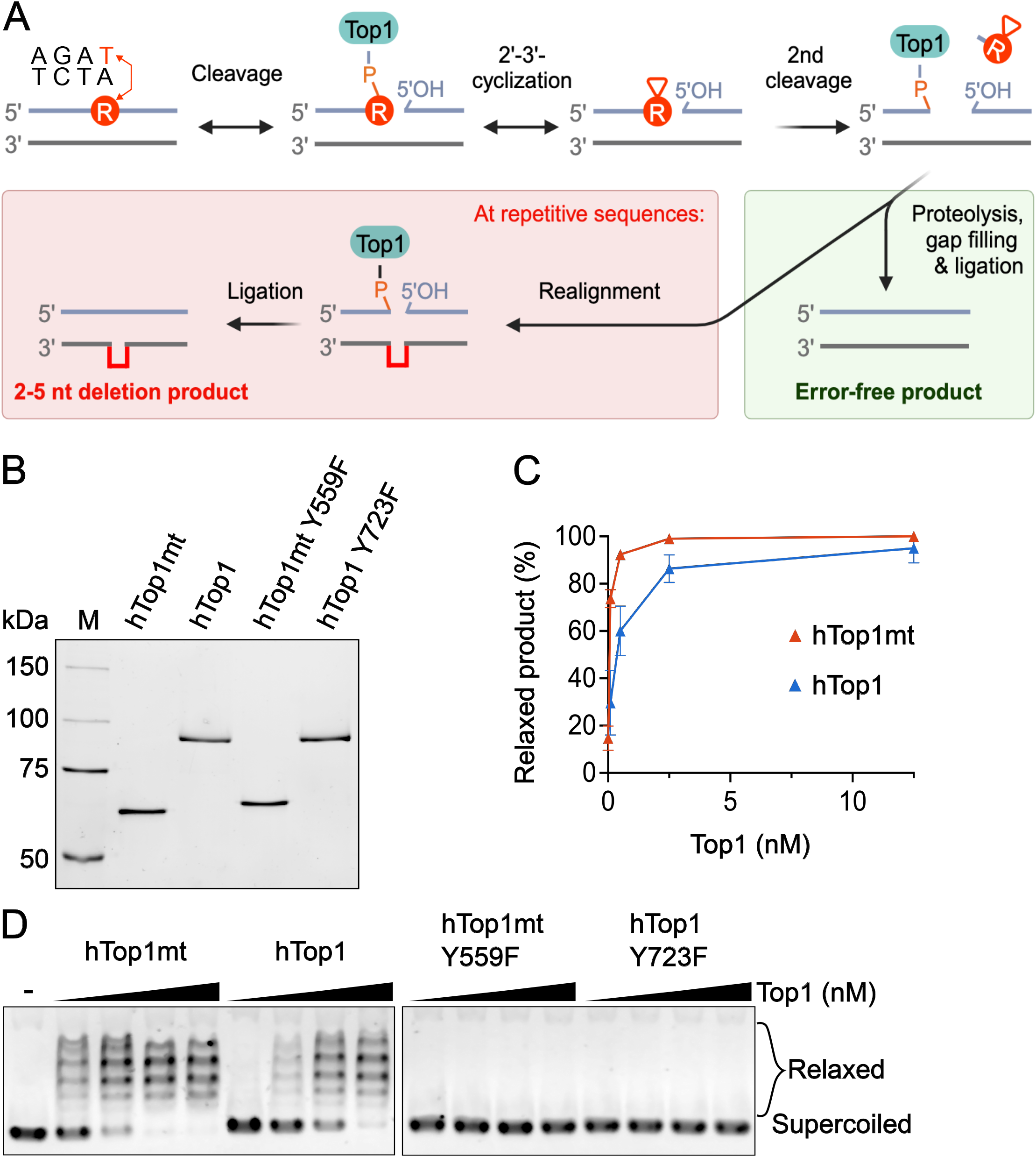
Relaxation of supercoiled DNA by purified hTop1mt and hTop1. **(A)** Model of Top1 action at an incorporated rNMP (*red circle*). Top1 cleavage at the rNMP generates a Top1-cc at the 3’ DNA end and a 5’-hydroxyl group. Attack by the 2’-OH of the rNMP on the 3’-phosphotyrosyl bond can prematurely release the Top1, creating a 2’,3’-cyclic phosphate that cannot undergo ligation without further processing. A second Top1 cleavage upstream of the cyclic phosphate can release a short DNA fragment containing the modified nucleotide, leaving a gapped intermediate with a covalently-associated Top1 that is too far from the 5’-OH for efficient ligation (*top row, far right*). An error-free product can be generated via proteolytic removal of the Top1, followed by gap filling and ligation (*green background*). Alternatively, strand realignment at a repetitive sequence can bring Top1 close enough to the 5’-OH to catalyze ligation of the two DNA ends, leading to deletion of one repeating unit of the repeat sequence (*red background; deleted region shown as red line*). **(B)** SDS-PAGE analysis of the wildtype and catalytically-dead variants of hTop1mt and hTop1. **(C)** Relaxation reactions containing 350 ng of pUC19 and increasing concentrations (0.1, 0.5, 2.5 and 12.5 nM) of wildtype (*left panel*) or catalytically-inactive (*right panel*) hTop1mt and hTop1 enzymes. **(D)** Quantification of the relaxation activity of the wildtype topoisomerases in Fig. 1C. The amount of relaxed product was quantified and expressed in percent of the total signal intensity in the lane. The average of three independent experiments is shown, and the error bars represent the standard error of the mean.

Like scTop1, the human nuclear hTop1 possesses endoribonuclease activity that results in short deletions at repetitive sequences (Sekiguchi and Shuman 1997; Zimmermann et al. 2018). However, whether the mitochondrial enzyme hTop1mt contains endoribonuclease activity has not been assessed. In this work, we sought to compare the endoribonuclease activities of the purified human nuclear and mitochondrial Top1s and found the mitochondrial enzyme to be less active on rNMPs than its nuclear paralog. This difference in ribonuclease activity was accordingly reflected in the enzymes’ ability to generate short deletions at repetitive sequences. These findings help explain why rNMPs are tolerated in the human mitochondrial genome.

## Materials and methods

### Overexpression and purification of recombinant protein

Wildtype and catalytically inactive variants of hTop1mt (excluding the mitochondrial targeting sequence), hTop1 and scTop1 were cloned into pRS424-GAL-GST where a recognition sequence for the rhinoviral 3C protease separates the GST tag from the N-terminus of the Top1 gene (Bylund, Majka, and Burgers 2006). Recombinant Top1 variants were expressed in *S. cerevisiae* strain PY116 (*MATa his 3–11,15 leu2-3,112 ura 3–52 trp1-Δ pep4–3 prb1–1122 prc1– 407 nuc1::LEU2*; (Chilkova, Jonsson, and Johansson 2003)); cell lysis and ammonium sulfate precipitation (0.31 g/ml) were essentially as previously described (Bylund, Majka, and Burgers 2006), with the following modifications: cells were grown in a LEX-48 bioreactor (Epiphyte 3), and lysed in the 6875 Freezer/Mill (SPEX) using 7 cycles with the following settings: T1= 2 min, T2= 2 min, T3= 5 min; impact frequency rate: 12 times/sec.

Ammonium-sulfate precipitated protein was resuspended in buffer HEP-0 (50 mM HEPES-NaOH pH 7.4, 10% glycerol, 1 mM dithiothreitol (DTT), 1 mM EDTA, 0.01% NP-40) and buffer added until the conductivity of the lysate equaled that of HEP-400 (HEP-0 with 400 mM NaCl). The lysate was added to equilibrated glutathione-Sepharose 4B beads (GE Healthcare; 1 ml/100 g cells) and gently rotated for 4 h at 4°C. The beads were collected at 500 × g in a swinging-bucket rotor, followed by batch washes (2 × 20 ml of HEP-400). The beads were transferred to a 10-ml column and washed with 40 ml of HEP-400 containing 1 mM pepstatin A, 1 mM AEBSF-HCL, 0.04 mM Aprotinin, 0.7 mM E-64 and 1.1 mM Leupeptin (Serva), then with 25 ml HEP-400 containing 5 mM MgCl_2_ and 1 mM ATP. This was followed by a wash with HEP-400 and then HEP-200. Protein was eluted in HEP-150 containing 30 mM reduced glutathione (pH adjusted to 8.1). Fractions that contained Top1 based on SDS-PAGE analysis were pooled and treated with 30 U rhinoviral 3C protease overnight at 4°C.

The following day, the Top1 protein was loaded onto a heparin column equilibrated in HEP-150 without protease inhibitors. The protein was washed with 10 column volumes of HEP-300 at a flow rate of 1.0 ml/min, and eluted in HEP-750 at a flow rate of 0.5 ml/min. Fractions containing pure protein were collected and dialyzed overnight against HEP-200 in a 10 000 MWCO Slide-A-Lyzer Dialysis cassette (ThermoScientific). Protein concentration was determined by densitometric analysis from SDS-PAGE gels containing a standard curve of bovine serum albumin (BSA; ThermoScientific).

### Preparation of dsDNA substrates

Oligonucleotides were purchased from Sigma Aldrich, IDT or Eurofins and are listed in Table S1. One nmol of each oligonucleotide was annealed to an equimolar amount of its complementary strand by denaturing at 95°C for 5 min in TE (50 mM Tris-HCl pH 8.0, 1 mM EDTA) containing 100 mM NaCl, and allowing the reaction mixture to cool to room temperature. The DNA was separated on a 15% acrylamide gel in 0.5 × TBE (15 mM Tris, 44.5 mM boric acid, 1 mM EDTA), stained with 3 × GelRed (Biotium) for 30 min and visualized by using Chemidoc™ (Bio-Rad). The bands corresponding to double-stranded molecules were excised, eluted from crushed gel slices into TE buffer (10 mM Tris-HCl, pH 8.0, 1 mM EDTA), and purified by phenol-chloroform extraction and isopropanol precipitation.

### DNA relaxation assay

The standard cleavage assay mixture contained 350 ng of supercoiled pUC19, 10 mM Tris-HCl pH 8.0, 50 mM KCl, 5 mM MgCl_2_, 0.1 mM EDTA, 5 mM DTT, 15 mg/ml BSA and the indicated concentration of Top1 enzymes. Reactions were performed at 30°C for 10 min and terminated by addition of 0.4% SDS and 6 × loading buffer (Thermo Scientific). The reaction mixture was loaded onto an 0.8% agarose gel in 1x TAE (40 mM Tris, 1 mM EDTA, 20 mM acetic acid) and separated at 120 V for 2 h. After electrophoresis, DNA bands were stained with 3 × GelRed and visualized on the Chemidoc imager (Bio-Rad).

### Cleavage assay

The standard 10 μl assay mixture contained 20 mM Tris-HCl pH 7.8, 5 mM MgCl_2_, 50 mM NaCl, 100 mg/ml BSA, 1 mM DTT, 10 mM DMSO, 50 nM Cy5-labeled dsDNA substrate and the indicated concentrations of Top1 in presence or absence of 10 μM camptothecin (CPT). Reactions were carried out at 30°C for 60 min and terminated by addition of 5 ml of 5 × Stop buffer (10 mM EDTA, 0.05% SDS) and 10 μl of formamide (40%). Samples were resolved on a 17% PAGE gel containing 7M urea in 1 × TBE at 60 W for 2.5 h. Fluorescence signal was detected on a Typhoon Biomolecular imager (Amersham) and quantified using ImageJ software (Java).

### Fluorescence anisotropy

Fluorescence anisotropy reactions containing 0.5 nM double-stranded oligonucleotides with a 5’-FAM labelled top strand and 0 – 60 nM Top1 in binding buffer (20 mM Tris pH 8, 66 mM NaCl, 1 mM DTT, 10% glycerol) were pipetted in triplicate onto black shallow 384-well microplates (OptiPlate-F, PerkinElmer) and incubated in the dark for 10 min at 25°C. Fluorescence intensities were measured from above on a CLARIOstar *Plus* plate reader (BMG Labtech) with the excitation and emission wavelengths 480 nm and 520 nm, respectively. Fluorescence anisotropy in millianisotropy units (mA) was calculated using MARS Data analysis Software (BMG Labtech) according to Equation 1: fluorescence anisotropy = (F - F)/(F +2* F)*1000, where F and F are the parallel and perpendicular emission intensity measurements corrected for background (buffer). The grating factor G was calculated using Equation 2: G = F / F and was equal to 1. The dissociation constant (K_d_) was determined by fitting the data to a quadratic equation by non-linear regression analysis in GraphPad Prism software (GraphPad Software, Inc., USA) using Equation 2:

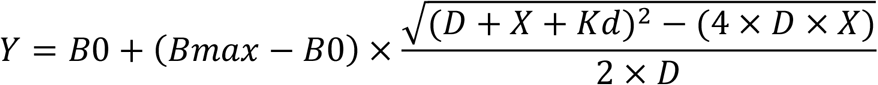

where Y is the anisotropy value at protein concentration X, X is the concentration of Top1 in nM, B0 and Bmax are specific anisotropy values associated with free DNA and the DNA-Top1 complex, respectively, and D is the concentration of DNA in nM. Reactions with different substrate concentrations yielded K_d_ values that did not significantly differ with those in Table S2 with 0.5 nM substrate, confirming the K_d_ values were not artefacts of analysis in the intermediate regime (Jarmoskaite et al. 2020).

## Results

### hTop1mt exhibits endoribonuclease activity

To compare the properties of hTop1mt and hTop1, we expressed and purified the recombinant wildtype (WT) proteins along with their catalytically-inactive variants where the catalytic tyrosine is mutated to a phenylalanine (Fig. 1b). Both WT enzymes showed robust activity on negatively supercoiled pUC19, with hTop1mt catalyzing full relaxation of the substrate at a lower enzyme concentration than hTop1 (Fig. 1c-d). As expected, the catalytically-inactive variants were unable to relax the substrate.

We next analyzed the ability of the Top1s to cleave a linear double-stranded 30-bp substrate containing the perfect consensus cleavage motif 5’-AGA**T**^**↓**^-3’ from the (AT)_2_ hotspot of the *S. cerevisiae CAN1* gene that exhibits a strong Top1-dependent deletion signature (Takahashi et al. 2011; Kim et al. 2011). Figure 2a shows the top strand of the double-stranded substrate, while the bottom strand is not shown for clarity. The experiments were carried out on a pair of substrates where the top strand consisted of all deoxyribonucleotides (substrate A) or where the deoxy-T at the cleavage site was replaced with a ribo-U (substrate B; rU in red in Fig. 2a). Substrates were fluorescently labelled on the 3’-end of the top strand, whereby cleavage at the preferred cleavage site should yield an 18-nucleotide product that can be detected if the religation step of the reaction is prevented either by addition of CPT or through the premature liberation of the enzyme upon 2’-3’-cyclization when the cleavage occurs at an rNMP (Fig. 1a).

**Figure 2.**
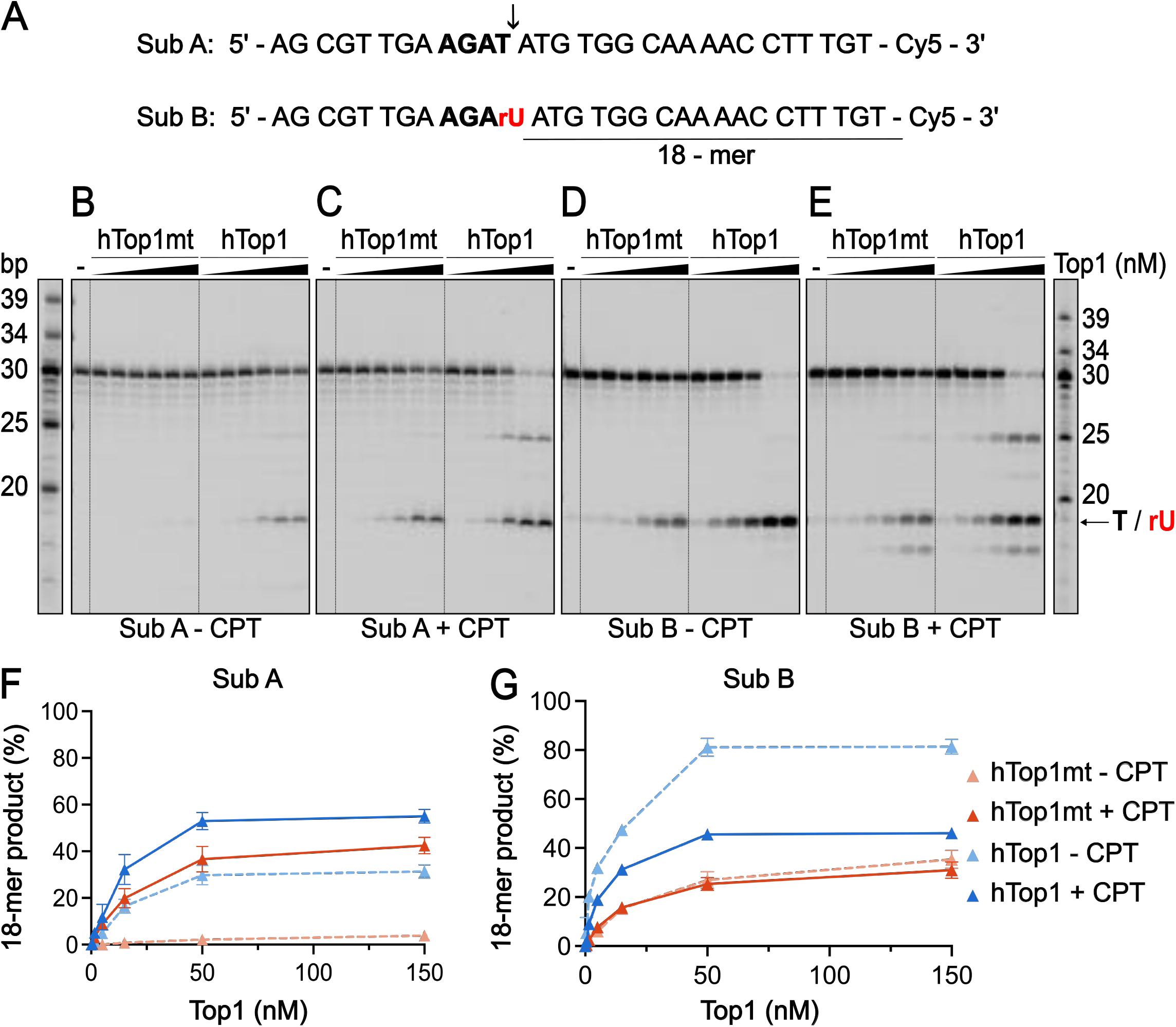
hTop1mt exhibits endoribonuclease activity. **(A)** The top strand of the dsDNA substrates containing the preferred cleavage motif *(bold)* found at the (AT)_2_ hotspot of the *S. cerevisiae CAN1* locus. The consensus cleavage site is marked by a *black arrow* and the resulting 18-nt product is *underlined*. The top strand was labelled with Cy5 at the 3’-end. Substrate B contains a rUMP (*red*) at the cleavage site. **(B-C)** Representative Top1 cleavage assays containing 50 nM substrate A and increasing concentrations (0.15, 0.5, 5, 15, 50 and 150 nM) of wildtype hTop1mt and hTop1 enzymes in the absence **(B)** and presence **(C)** of 10 μM camptothecin *(CPT)*, an inhibitor of the re-ligation step. **(D-E)** Cleavage assays on substrate B with an rUMP at the cleavage site in the absence **(D)** and presence **(E)** of CPT. **(F-G)** Quantification of the 18-mer cleavage product from assays on substrate A **(F)** and substrate B **(G)**. The amount of 18-mer product was quantified and expressed in percent of the total signal intensity in the lane. The average of the three independent experiments is shown, and the error bars represent the standard error of the mean.

Accordingly, addition of CPT clearly increased the intensity of the product bands, including that of the main 18-nt product, in reactions containing the all-DNA substrate A (Fig. 2, compare panels b and c). On substrate B containing a rUMP at the cleavage site, frequent cleavage events at the ribonucleotide were readily detected even in the absence of CPT, confirming they occurred at the rUMP (Fig. 2d). Addition of the inhibitor revealed the presence of additional, weaker cleavage sites, one of which was located a few nucleotides downstream of the rUMP (Fig. 2, compare panels d and e), well in line with the reported preference of Top1 to cleave 2-6 nt from a nick (Christiansen and Westergaard 1999). The results of Fig. 2 demonstrate that like its nuclear homolog, the human mitochondrial Top1 exhibits endoribonuclease activity. However, cleavage by nuclear hTop1 was somewhat more efficient than that by hTop1mt on these linear dsDNA substrates both with and without an rNMP at the cleavage site (Fig. 2f-g, compare solid blue to solid red line).

### hTop1mt and hTop1 differ in their propensity to act at an upstream rNMP

Next, we analyzed the preference of the Top1s for cleavage at an rNMP over a dNMP. To this end, we used a 34-mer dsDNA substrate derived from another *CAN1* deletion hotspot, the (TC)_3_ that has been thoroughly studied in the context of Top1-mediated rNMP repair (Kim et al. 2011; Sparks and Burgers 2015). The activity of the two Top1s was compared on the all-DNA substrate (substrate C) and on substrate D that contained a single rNMP located 2 nt upstream of the preferred cleavage site (Fig. 3a). On this pair of substrates, cleavage at the perfect consensus cleavage site should generate a 16-mer product, while cleavage at the upstream rGMP should yield an 18-mer. The action of both Top1s on the all-DNA substrate C yielded only the 16-mer product, indicating that this substrate contained only the one site corresponding to the consensus cleavage motif (Fig. 3b-c). However, a longer 18-nt product formed through cleavage at the upstream rGMP was observed in reactions containing the ribonucleotide-containing substrate D both in the absence and presence of CPT (Fig. 3d-e). Interestingly, the 18-mer product was only generated by hTop1 and not by hTop1mt (Fig. 3h-i). These results indicate that an embedded rNMP is able to attract hTop1 to cleave beyond its consensus cleavage site, while hTop1mt acts only at the consensus site. As seen by the appearance of a 32-mer deletion product in reactions with hTop1 in Fig. 3d, this behavior has implications for Top1-dependent deletion formation that will be more closely addressed in Fig. 5.

**Figure 3.**
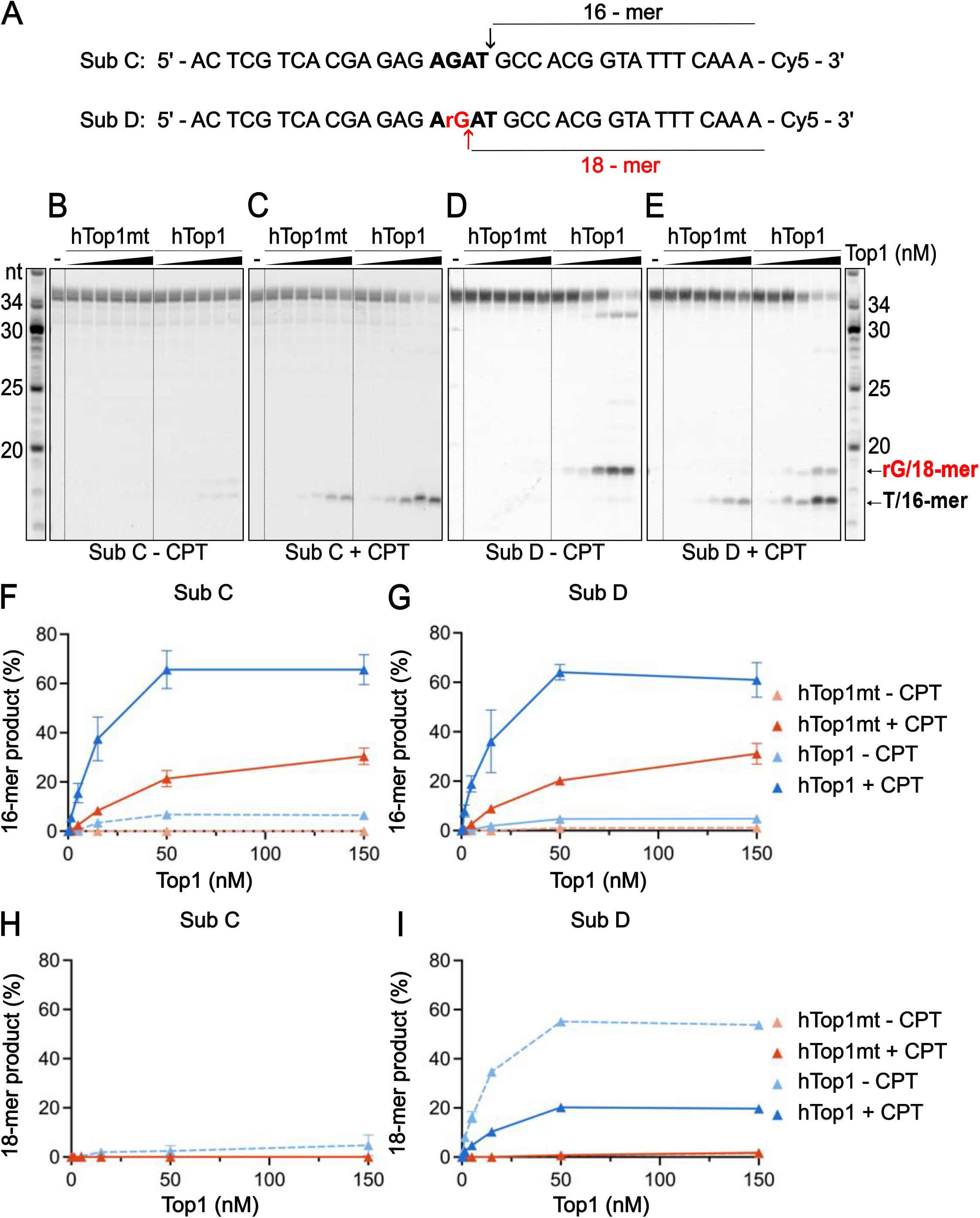
hTop1mt and hTop1 differ in their propensity to cleave at an rNMP beyond the preferred cleavage site. **(A)** The top strand of the dsDNA substrates containing the preferred cleavage motif *(bold)* modified from the (TC)_3_ hotspot of the *S. cerevisiae CAN1* locus. The top strand was labelled with Cy5 at the 3’-end. The cleavage site is marked by a *black arrow* and the resulting 16-nt product is indicated. Substrate D contains an rGMP *(red)* 2 nt upstream of the consensus cleavage site; cleavage at the rGMP generates an 18-nt product *(red arrow)*. **(B-C)** Representative Top1 cleavage assays containing 50 nM substrate C and increasing concentrations (0.15, 0.5, 5, 15, 50 and 150 nM) of wildtype hTop1mt and hTop1 enzymes in the absence **(B)** and presence **(C)** of 10 μM camptothecin *(CPT)*. **(D-E)** Cleavage assays in the absence **(D)** and presence **(E)** of CPT were carried out as in Fig. 3B-C but on substrate D with an rGMP upstream of the cleavage site. **(F-G)** Quantification of the 16-mer cleavage product from assays on substrate C **(F)** and substrate D **(G). (H-I)** Quantification of the 18-mer cleavage product from assays on substrate C **(H)** and substrate D **(I)**. The amount of 18-mer and 16-mer products were quantified and expressed in percent of the total signal intensity in the lane. The average of three independent experiments is shown, and the error bars represent the standard error of the mean.

### The DNA binding behavior of hTop1mt on rNMP-containing substrates differs from its homologs

We next investigated the impact of a single rNMP on the DNA binding affinity of the Top1 enzymes using fluorescence anisotropy. In addition to the two human Top1s, these experiments were expanded to include their *Saccharomyces cerevisiae* homolog (scTop1) that is dually-localized to both the nuclear and mitochondrial compartments (Wang et al. 1995) (see Fig. S1 for SDS-PAGE analysis and DNA relaxation activity of scTop1). To avoid confounding effects due to differences in enzyme activity and thus the efficiency of covalent complex formation during the enzymes’ reaction cycle, we limited our analysis to noncovalent binding by using the catalytically-inactive Top1 variants, a strategy successfully employed in studies of *e*.*g*. the related vaccinia virus Top1 (Sekiguchi and Shuman 1994). Binding was investigated on the two dsDNA substrate pairs used in Figs. 2 and 3: substrates A *vs*. B where the rNMP is located at the cleavage site (AGA<u>T</u>^↓^ *vs*. AGA<u>rU</u>^↓^), and substrates C *vs*. D with the rNMP located upstream of the cleavage site (A<u>G</u>AT^↓^ *vs*. A<u>rG</u>AT^↓^) (Fig. 4A, D). The anisotropy values for each Top1 complexed with each of the four substrates (Fig. 4b-c, e-f) were used to determine their respective K_d_ values (Fig. 4g-i, exact values in Table S2).

**Figure 4.**
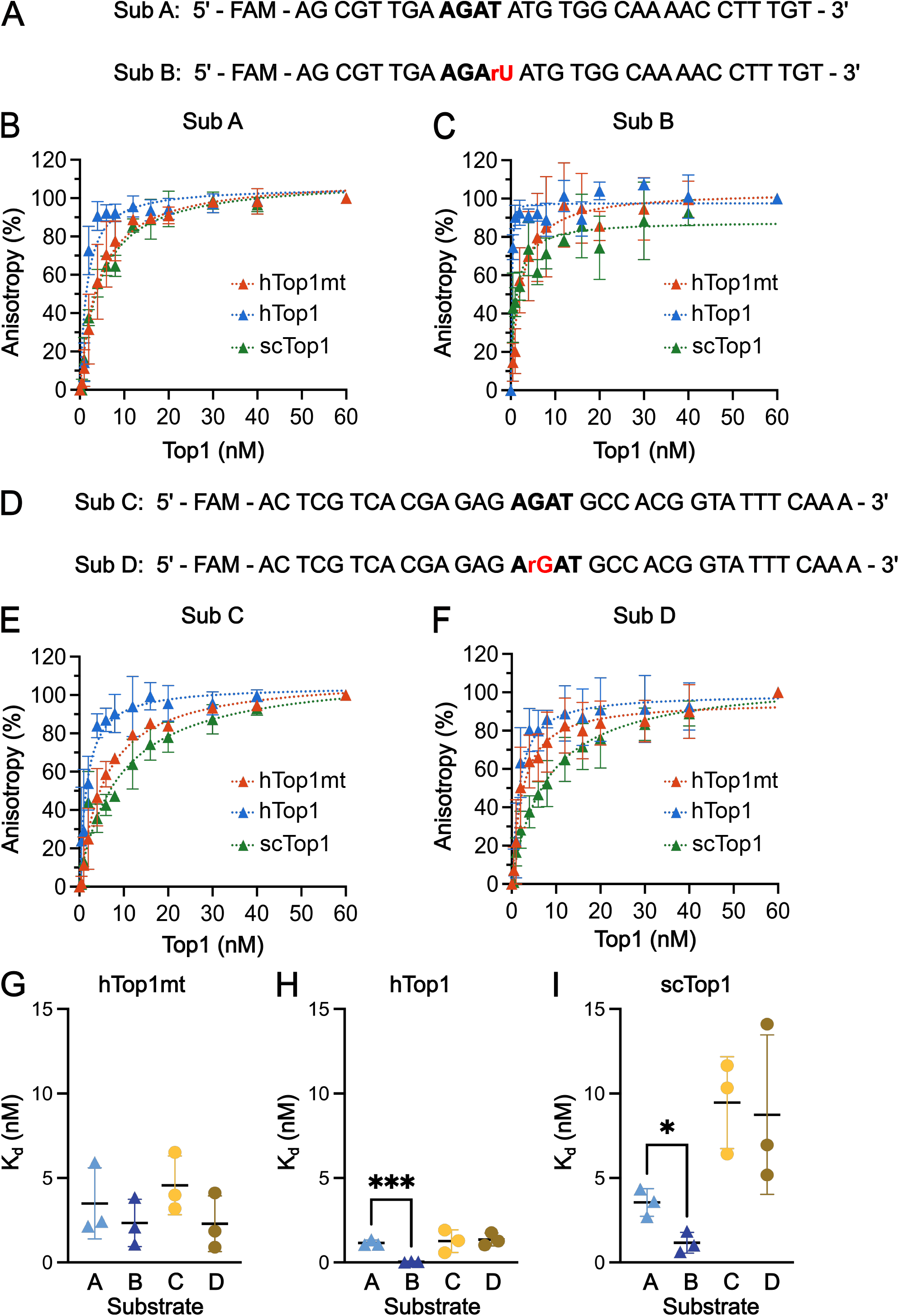
hTop1mt shows lower affinity for a substrate containing an rNMP at the cleavage site. **(A)** The sequences of the 5’-FAM labelled top strands of the dsDNA substrates used for the fluorescence anisotropy measurements in Fig. 4B-C. **(B-C)** Quantification of the DNA binding affinity of the Top1s by fluorescence anisotropy. The binding reaction contained 0.5 nM substrate A **(B)** or substrate B **(C)** along with increasing concentrations (0.5, 1, 2, 4, 6, 8, 12, 16, 20, 30, 40, 60 nM) of catalytically-dead Top1 variants (hTop1mt-Y559F, hTop1-Y723F and the *Saccharomyces cerevisiae* scTop1-Y272F). **(D)** The sequences of the 5’-FAM labelled top strands of the dsDNA substrates used for the fluorescence anisotropy measurements in Fig. 4E-F. **(E-F)** Quantification of the DNA binding affinity of the Top1s performed as in Fig. 4B-C but with 0.5 nM substrate C **(E)** or substrate D **(F)**. The average of three independent experiments is shown, and the error bars represent the standard error of the mean. **(G-I)** The K_d_ values of hTop1mt **(G)**, hTop1 **(H)** and scTop1 **(I)** for each of the four substrates A-D was determined based on the anisotropy data in Fig. 4b-c, e-f using the quadratic equation. The mean of three independent experiments is shown, error bars represent the standard deviation. Pairs of substrates were compared using t-tests; *<0.05, ***<0.001. The exact values are presented in Table S2.

**Figure 5.**
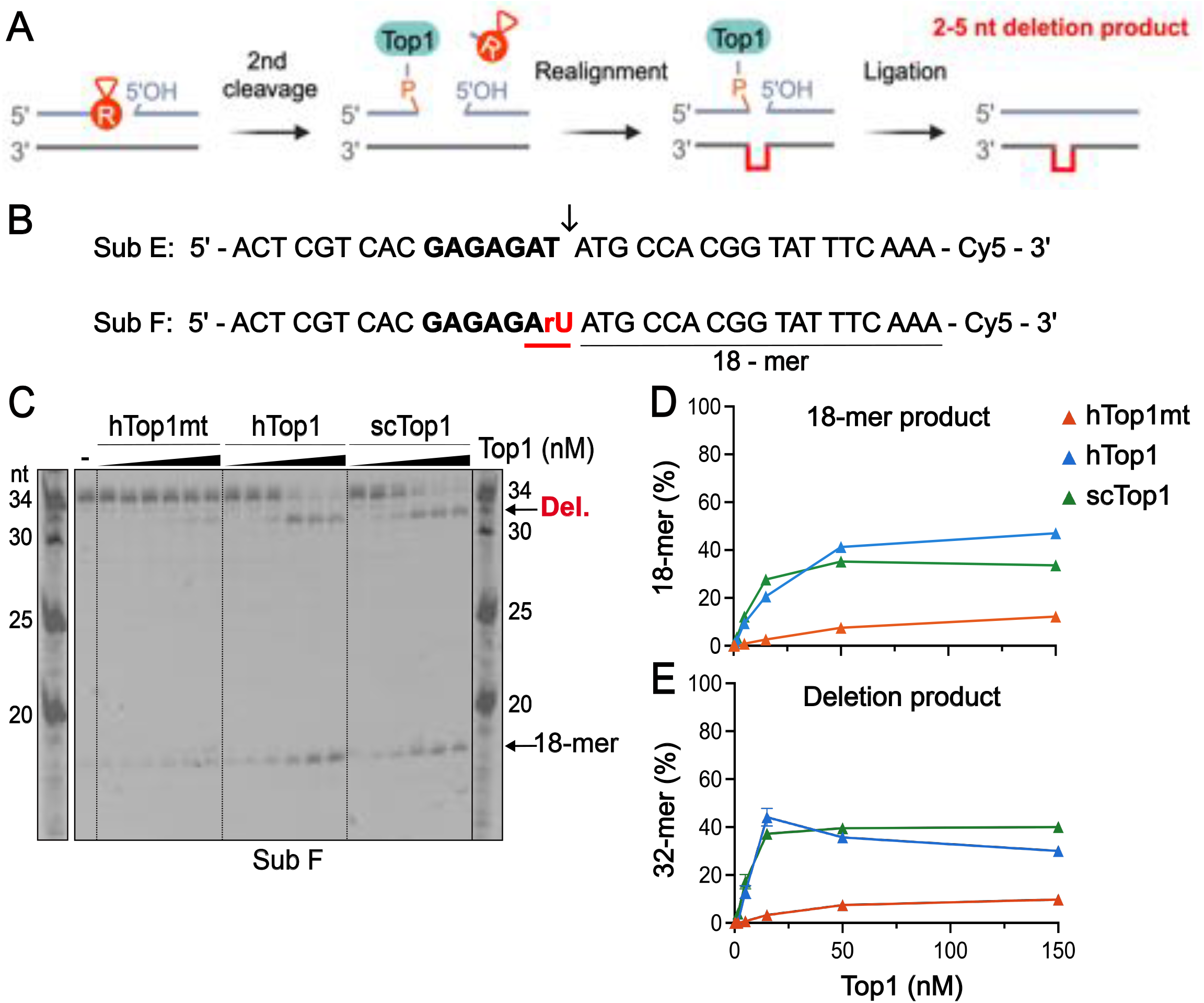
hTop1mt-dependent deletion formation is a rare event. **(A)** Schematic representation of deletion formation on a dsDNA substrate containing an rNMP at a repetitive sequence in the top strand. The Top1-cc at the rNMP is dissolved following the formation of a 2’,3’-cyclic phosphate (*not shown*). A second Top1 cleavage a few nt upstream releases the fragment with the cyclic phosphate. Realignment of the two strands relative to each other extrudes one repeating unit in the repeat sequence (*red line*), bringing the 3’-phosphate-Top1 close enough to the 5’-hydroxyl end for Top1 to finish its reaction cycle through ligation of the DNA ends. This results in deletion of one repeat unit in the top strand. **(B)** The top strand of the dsDNA substrates E and F containing the preferred cleavage motif *(bold)* modified from the (TC)_3_ hotspot of the *S. cerevisiae CAN1* locus. The top strand was labelled with Cy5 at the 3’-end. The cleavage site is marked by a *black arrow* and the resulting 18-nt product is indicated. Substrate F contains a rUMP *(red)* at the consensus cleavage site, cleavage at which can result in a 2-nt deletion *(red underline)*. **(C)** Representative Top1 cleavage assay containing 50 nM substrate F and increasing concentrations (0.15, 0.5, 5, 15, 50 and 150 nM) of wildtype hTop1mt, hTop1 and scTop1 enzymes in the absence of CPT. **(D)** Quantification of the 18-mer cleavage product in the reactions in Fig. 5c. **(E)** Quantification of the 32-mer deletion product in the reactions in Fig. 5c. The amount of the 18-mer and 32-mer products was quantified and expressed in percent of the total signal intensity in the lane. The average of the three independent experiments is shown, and the error bars represent the standard error of the mean.

A first comparison of the K_d_ values on the two all-DNA substrates A and C indicates that hTop1mt exhibits somewhat lower affinity to both all-DNA substrates than does hTop1 (Fig. 4g *vs*. h, sub A and sub C; Table S2). A similar conclusion has previously been drawn based on the chromatin retainment of hybrid Top1 enzymes *in cellulo* (Dalla Rosa et al. 2009). This lower DNA binding affinity likely contributes to the somewhat lower cleavage efficiency of hTop1mt on linear DNA substrates (Fig. 2f and Fig. 3f).

We then analyzed the effect of a single rNMP on substrate binding. The replacement of the T^↓^ at the cleavage site with rU^↓^ increased the affinity of both hTop1 and scTop1 for the DNA substrate, while the binding affinity of hTop1mt was not impacted by the presence or absence of the rU (Fig. 4g-i, substrate A *vs*. B). In contrast, the inclusion of an rNMP at a position 2 nt upstream of the cleavage site had little effect on any of the enzymes’ affinity for the DNA substrate (Fig. 4g-i, substrate C *vs*. D). These findings reveal a differential impact of an incorporated rNMP on the binding affinity of hTop1 and scTop1 on one hand and hTop1mt on the other: the precise location of the rNMP within the consensus motif affects the strength of binding by hTop1 and scTop1, resulting in the observed higher affinity only if the rNMP is located precisely at the cleavage site. Conversely, binding by hTop1mt is not appreciably affected by an rNMP, regardless of the location of the rNMP. In conclusion, the binding behavior of hTop1mt to rNMP-containing substrates differs from what is observed for hTop1 and scTop1, both of which show significantly higher affinity to a DNA substrate with an rNMP in the cleavage site.

### In contrast to hTop1 and scTop1, hTop1mt action rarely leads to rNMP-dependent deletions

As mentioned above, the action of hTop1 and scTop1 at rNMPs has been shown to promote the generation of 2-5 nt deletions when the rNMP is present at a repetitive sequence that allows strand slippage to occur (Kim et al. 2011; Sparks and Burgers 2015; Sekiguchi and Shuman 1997). Briefly, deletions arise through a mechanism that involves a second Top1 cleavage event a few nt upstream of the 2’,3’-cyclic phosphate, followed by strand slippage that extrudes one repeating unit of the repeat sequence. This strand realignment brings the Top1-bound 3’-phosphate end of the DNA close enough to the 5’-hydroxyl end for the enzyme to catalyze ligation of the two ends (Fig. 5a). As a result, one repeating unit of the repeat sequence is deleted from the top strand, and replication of this strand will yield a dsDNA molecule lacking one repeat unit from both strands.

Our results so far indicate that compared to hTop1, hTop1mt shows less preference for binding a DNA substrate with an rNMP at the cleavage site and incises somewhat less efficiently at a cleavage-site rNMP (Table S2; Fig. 2b). We next asked how efficiently hTop1mt mediates rNMP-dependent deletion formation at repetitive sequences. For this, the sequence of the *CAN1* (TC)_3_ deletion hotspot was modified by insertion of an AT dinucleotide immediately 3’ of the cleavage site, an alteration that is known to increase the efficiency of deletion formation (Sparks and Burgers 2015). The resultant substrates E and F contained the consensus T or a rU, respectively, at the cleavage site (Fig. 5b).

The action of hTop1mt, hTop1 and scTop1 on the rUMP-containing substrate F yielded the expected 18-nt cleavage product as well as a 32-mer deletion product (Fig. 5c-e). However, in line with the relatively inefficient cleavage at the rUMP observed with hTop1mt, the yield of deletion product generated by hTop1mt was significantly lower than with hTop1 or scTop1: up to 9.7%, 30% and 40% of the substrate was converted to the 32-mer deletion product by hTop1mt, hTop1 and scTop1, respectively (Fig. 5d, e). As expected for an rNMP-specific deletion mechanism, no deletion formation was observed on the all-DNA substrate E (Fig. S2a). Instead, Top1 action on substrate E yielded only the 18-mer product that corresponds to cleavage at the consensus cleavage site, and this product was only observed in the presence of CPT (Fig. S2b). Of note is that in the presence of CPT, hTop1mt action on the all-DNA substrate E generated more of the 18-mer cleavage product than did hTop1 (Fig. S2b, c). These results indicate that while hTop1mt efficiently cleaves at the cleavage motif in the absence of an rNMP, the presence of a ribonucleotide at the cleavage site lowers its cleavage efficiency in comparison to hTop1 and scTop1. Consequently, hTop1mt-mediated deletion formation is a rare event.

## Discussion

Previous work in the field of mtDNA maintenance has uncovered a differential impact of the physiological level of rNMPs on mtDNA stability in yeast and mammalian cells — while a decrease in rNMP frequency had a positive impact on mtDNA stability in *S. cerevisiae*, it had no observable effect on the integrity of mouse mtDNA (Wanrooij et al. 2017; Wanrooij et al. 2020). The different properties of the mitochondria-resident Top1s illuminated in the current study offer at least one contributing mechanism to explain the observed discrepancy between organisms. We find that the human mitochondrial hTop1mt lacks the binding preference that the nuclear human hTop1 and the *S. cerevisiae* scTop1 demonstrate for substrates with an rNMP at the cleavage site (Fig. 4; Table S2). Top1mt also shows somewhat lower cleavage efficiency at rNMPs than its nuclear counterpart, to an extent that differs from substrate to substrate (Fig. 2, Fig. 5). This results in a 4-fold lower efficiency of deletion formation compared to hTop1 on the modified *CAN1* substrate utilized in Fig. 5.

Most strikingly, while hTop1 can shift to cleave at an rNMP located outside of the consensus cleavage site, the same behavior is not observed for hTop1mt (Fig. 3). Like hTop1, scTop1 showed efficient cleavage at the rG upstream of the cleavage site on substrate D (Fig. S2d-f). This differing feature between hTop1mt and the other two Top1s is expected to have a major impact *in cellulo*, where the substrate of hTop1mt is an mtDNA molecule with a random distribution of incorporated rNMPs, most of which will be located outside of a Top1 cleavage site. Mammalian mtDNA contains ca 6-31 incorporated rNMPs per strand, which corresponds to an average frequency of one rNMP per 530-2700 nts (Berglund et al. 2017; Forslund et al. 2018; Wanrooij et al. 2020). It is thus very unlikely to find an incorporated rNMP precisely at a Top1 consensus cleavage site, and there will therefore be very few opportunities for hTop1mt to act at an rNMP. The likelihood is decreased further by the fact that rU, that should be the rNMP incorporated at the consensus cleavage site 5’-AGA**T**^**↓**^-3’, is the least frequent rNMP found in mammalian mtDNA, both in mouse and human cells (Wanrooij et al. 2020; Berglund et al. 2017). However, based on the findings in Fig. 3, the properties of hTop1 and scTop1 would allow them to act even at an rNMP outside the cleavage site and thus potentially generate deletions and/or nicked repair intermediates at far more positions than hTop1mt. Taken together, these properties of hTop1mt can partially explain why rNMPs have less of a negative impact on mtDNA in mammalian compared to yeast cells.

A comparison can also be made with nuclear DNA, where the presence of rNMPs has been well-documented to lead to both short Top1-mediated deletions and other signs of genome instability (Kim et al. 2011; Williams et al. 2013; Zimmermann et al. 2018). The contrasting tolerance of rNMPs in mtDNA can be explained by at least three features of this genome that differ from the nuclear one. First, the considerable redundancy inferred by the multicopy nature of the mitochondrial genome makes it less vulnerable upon induction of strand breaks by rNMPs, as damage to one mtDNA molecule would still leave the cell with many intact copies. Should the reactivity of rNMPs result in a double-strand break, the damaged mtDNA molecule can be cleared by the exonuclease activity of the mitochondrial replicative polymerase Pol γ and by the MGME1 nuclease (Peeva et al. 2018). A second reason for rNMP tolerance in the mtDNA is the reverse transcriptase activity of Pol γ that allows it to read through a template-strand rNMP far more efficiently than nuclear replicative polymerases, resulting in fewer termination events (Forslund et al. 2018; Kasiviswanathan and Copeland 2011; Göksenin et al. 2012; Clausen et al. 2013; Watt et al. 2011). The third contributing factor to the tolerance of rNMPs in the mtDNA is the lower activity of hTop1mt at incorporated rNMPs uncovered in this study — thus, Top1-mediated rNMP repair is far less likely to generate adverse outcomes like deletions in the mtDNA than in the nuclear DNA. Interestingly, others have reported mtDNA loss following artificial targeting of hTop1 to human mitochondria (Dalla Rosa et al. 2009). However, the depletion was independent of hTop1 catalytic activity, and can therefore not be ascribed to cleavage at mtDNA rNMPs.

Our study did not address the activity of hTop1mt on mtDNA *in vivo*. It would be interesting to follow the consequences of targeting hTop1 or scTop1 to mitochondria in cells with a high or low mtDNA rNMP load. Another as-of-yet unanswered question is which residue(s) of hTop1 and hTop1mt contribute to the observed differences in their endoribonuclease activity. Theoretically, the mutation of the corresponding amino acids in hTop1mt could yield a mitochondrial topoisomerase 1 with an improved rNMP repair activity and a higher frequency of rNMP-dependent deletions in mtDNA. Further work is ongoing to answer these intriguing questions related to mtDNA stability that must be maintained in order to avoid disease.

## Acknowledgements

We warmly thank Prof. Erik Johansson and Dr. Göran Bylund for the pRS424-GAL-GST expression plasmid and the PY116 expression strain. Dr. Justin Sparks and Prof. Peter Burgers are gratefully acknowledged for insightful discussions, Dr. Igor Iashchishyn for expert guidance on fluorescence anisotropy, and Ms. Newal Jemal and Ms. Josephine Åberg for technical assistance. We acknowledge the Chemical Biology Consortium Sweden (CBCS) at Umeå University for access to the CLARIOstar *Plus* plate reader, and the Protein Expertise Platform at Umeå University (a part of Protein Production Sweden) for cloning assistance. This work was supported by grants from the Swedish Research Council (grant number 2019-01874), The Swedish Cancer Society (grant numbers 19 0022 JIA; 22 2381 Pj), The Knut and Alice Wallenberg Foundation (KAW 2021.0053), and the Kempe Foundations (JCK22-0016) to P.H.W.

## Author contributions

CPJB, EK, JMEF-investigation, methodology, formal analysis, validation, visualization; MW-investigation, validation; PHW-conceptualization, formal analysis, data curation, visualization, funding acquisition, project administration, supervision, writing-original draft. All authors reviewed and edited the final version.

## Declaration of competing interests

None.

## Notes

### Competing Interest Statement

The authors have declared no competing interest.

