## Supplemental data for "Biochemical analysis of the endoribonuclease activity of the human mitochondrial topoisomerase 1"

### Supplementary tables

#### Table S1. Oligonucleotides used in this study

The Top1 consensus motif is underlined; the identity and position of potential fluorescent labels is indicated in each respective figure.

| Name | Sequence |
|---|---|
| A | 5'-AGC GTT GAA GAT ATG TGG CAA AAC CTT TGT-3' |
| B | 5'-AGC GTT GAA GA**rU** ATG TGG CAA AAC CTT TGT-3' |
| Rev. comp. AB | 5'-ACA AAG GTT TTG CCA CAT ATC TTC AAC GCT-3' |
| C | 5'-ACT CGT CAC GAG AGA GAT GCC ACG GTA TTT CAA A-3' |
| D | 5'-ACT CGT CAC GAG AGA **rG**AT GCC ACG GTA TTT CAA A-3' |
| Rev. comp. CD | 5'-TTT GAA ATA CCG TGG CAT CTC TCT CGT GAC GAG T-3' |
| E | 5'-ACT CGT CAC GAG AGA **T**A TGC CAC GGT ATT TCA AA-3' |
| F | 5'-ACT CGT CAC GAG AGA **rU**A TGC CAC GGT ATT TCA AA-3' |
| Rev. comp. EF | 5'-TTT GAA ATA CCG TGG CAT ATC TCT CGT GAC GAG T-3' |

#### Table S2. DNA binding affinities of Top1 enzymes to substrates A-D

| $K_d$ (nM) | Sub A | Sub B | **Sub A *vs.* B** | Sub C | Sub D | **Sub C *vs.* D** |
|---|---|---|---|---|---|---|
| **hTop1mt** | 3.5 ± 2.1 | 2.34 ± 1.40 | ns | 4.57 ± 1.73 | 2.29 ± 1.65 | ns |
| **hTop1** | 1.16 ± 0.17 | 0.05 ± 0.02 | *** | 1.26 ± 0.67 | 1.36 ± 0.38 | ns |
| **scTop1** | 3.55 ± 0.82 | 1.17 ± 0.61 | * | 9.47 ± 2.72 | 8.75 ± 4.27 | ns |

**Supplementary figure legends**

**Supplementary Figure 1, related to Figure 4. The purity and relaxation activity of scTop1. (A)** SDS-PAGE analysis of the wildtype and catalytically-dead variants of the *Saccharomyces cerevisiae* Top1 (scTop1). **(B)** Relaxation reactions containing 350 ng of pUC19 and increasing concentrations (0.1, 0.5, 2.5 and 12.5 nM) of wildtype (*left panel*) or catalytically-inactive (*right panel*) scTop1 enzymes. **(C)** Quantification of the relaxation activity of the wildtype scTop1 from Fig. S1B, along with the wildtype hTop1mt and hTop1 from Fig. 1C. The amount of relaxed product was quantified and expressed in percent of the total signal intensity in the lane. The average of three independent experiments is shown, and the error bars represent the standard error of the mean.

**Supplementary Figure 2, related to Figures 3 and 5. No deletion formation on an all-DNA substrate, and scTop1 cleaves at an rG upstream of the cleavage site. (A**) Representative Top1 cleavage assay containing 50 nM substrate E and increasing concentrations (0.15, 0.5, 5, 15, 50 and 150 nM) of wildtype hTop1mt, hTop1 and scTop1 enzymes in the absence of CPT. **(B)** Representative Top1 cleavage assay containing 50 nM substrates E (*left panel*) or F (*right panel*) and 150 nM wildtype hTop1mt, hTop1 and scTop1 enzymes in the presence of 10 µM CPT. **(C)** Quantification of the 18-mer cleavage product in the reactions with CPT in Fig. S2b. **(D-E)** Representative Top1 cleavage assay on substrate C without an rNMP **(D)** or substrate D with an rG upstream of the cleavage site **(E)** and increasing concentrations (0.15, 0.5, 5, 15, 50 and 150 nM) of wildtype scTop1 in the absence of CPT. **(F)** Quantification of the 18-mer cleavage product in the reactions in Fig. S2d-e. The amount of the 18-mer product was quantified and expressed in percent of the total signal intensity in the lane. The average of the three independent experiments is shown, and the error bars represent the standard error of the mean.

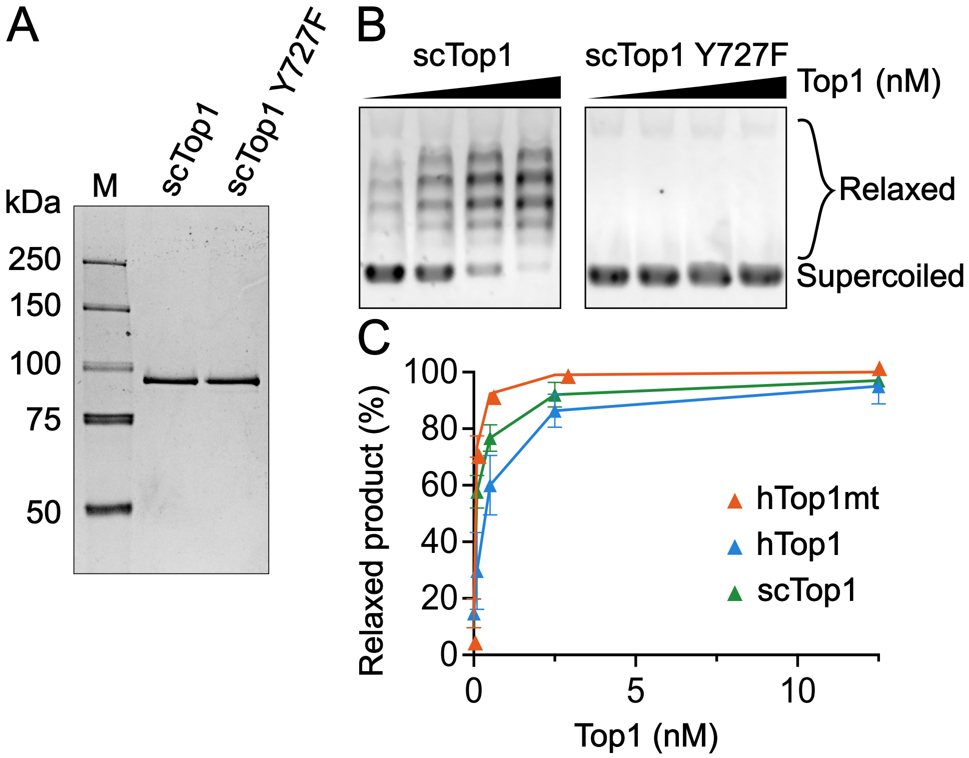
A
M
scTop1
scTop1 Y727F
kDa
250
150
100
75
50
B
scTop1
scTop1 Y727F
Top1 (nM)
Relaxed
Supercoiled
C
Relaxed product (%)
100
80
60
40
20
0
0
5
10
Top1 (nM)
hTop1mt
hTop1
scTop1

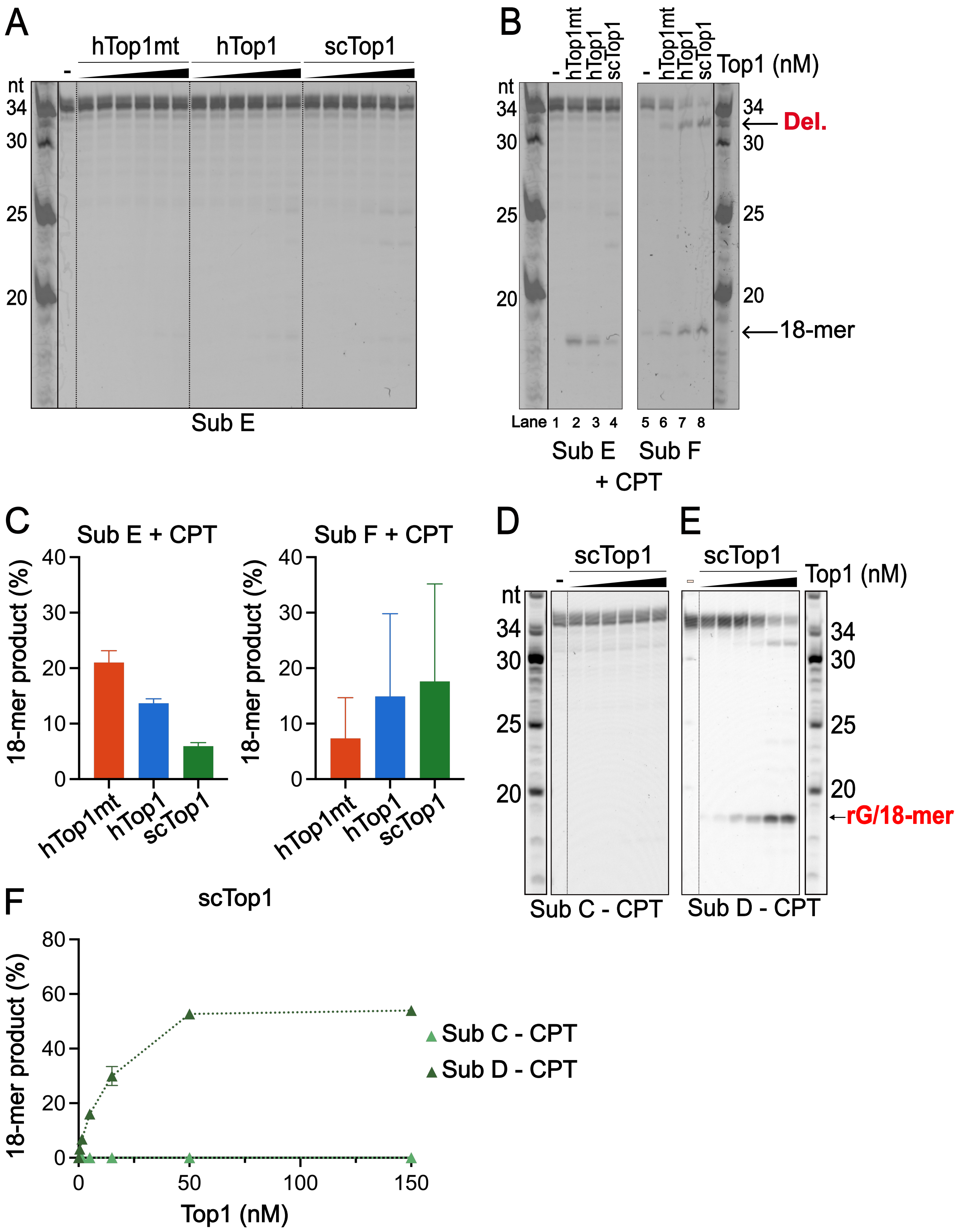
A
hTop1mt
hTop1
scTop1
nt
34
30
25
20
Sub E
B
hTop1mt
hTop1
scTop1
Top1 (nM)
Del.
18-mer
Lane 1 2 3 4 5 6 7 8
Sub E
Sub F
+ CPT
C
Sub E + CPT
Sub F + CPT
18-mer product (%)
hTop1mt
hTop1
scTop1
D
scTop1
Sub C - CPT
E
scTop1
Top1 (nM)
rG/18-mer
Sub D - CPT
F
scTop1
18-mer product (%)
Top1 (nM)
Sub C - CPT
Sub D - CPT
